# Antagonistic crosstalk fine-tunes sensitivities of IRE1 and PERK signaling during unfolded protein response

**DOI:** 10.1101/2020.08.09.243527

**Authors:** Jun Jiang, Yimiao Qu, Xiang Liu, Chao Tang, Ping Wei

## Abstract

When endoplasmic reticulum (ER) stress occurs, a collection of phylogenetically conserved signaling pathways, termed unfolded protein response (UPR) pathways, monitors the stress level in the ER and is activated to restore homeostasis. If stress is overwhelming, activation of these signaling pathways also leads to apoptosis. The initial response in the ER is fluxed into several parallel branches, i.e., IRE1, PERK, and ATF6 branch. How they coordinate in response to different ER stress levels remains largely unknown. Here, we constructed a dual-reporter system to simultaneously monitor and quantify the response of both the IRE1 and PERK branches. We found that the IRE1 and PERK branches were highly coordinated via mutual inhibition. Furthermore, IRE1 branch was more sensitive to ER stress than the PERK branch under low ER stress and IRE1 activity was attenuated under high ER stress. The differential sensitivity between the two branches arises from the interbranch inhibitor p58^IPK^, rather than the intra-branch inhibitor GADD34. Our results suggested a model where cells use the antagonistic crosstalk between parallel UPR signaling pathways to fine-tune their activities in response to different ER stress levels.

## Introduction

Mammalian cells sense ER stress levels *via* transducers on the ER membrane and activate downstream signaling cascades to restore protein folding homeostasis (Ron and Walter, 2007; Walter and Ron, 2011). When protein folding homeostasis is not restored, the UPR can induce apoptosis of the damaged cells, indicating that both the ER stress level and the UPR affect cell fate.

IRE1 and PERK branch, which are two main UPR signaling pathways, can induce both cytoprotective and proapoptotic responses (Ron and Walter, 2007; Walter and Ron, 2011). IRE1 and PERK are transducers on the ER membrane of IRE1 and PERK branch, respectively. Activated IRE1 specifically cleaves XBP1 mRNA, resulting in the translation of the bZIP transcription factor XBP1s, which induces cytoprotective responses by promoting the expression of stress response genes (Calfon et al., 2002; Jonikas et al., 2009; Lee et al., 2003). Activated IRE1 can also induce mRNA decay, decreasing the load of protein folding (Hollien et al., 2009; Hollien and Weissman, 2006). Activated PERK phosphorylates the translation initiation factor eIF2 on its α subunit (eIF2α), downregulating global protein translation to decrease the load of proteins in the ER (Harding et al., 1999). Phosphorylated eIF2α upregulates the translation of the transcription factor ATF4, which also induces stress response genes that restore protein folding homeostasis (Harding et al., 2000; Lu et al., 2004). When ER stress is overwhelming, PERK can induce proapoptotic effects mainly by CHOP (Puthalakath et al., 2007). However, CHOP also induces the expression of GADD34, which acts as a feedback inhibitor of the PERK branch by repressing eIF2α phosphorylation and represses cell apoptosis (Ma and Hendershot, 2003; Marciniak et al., 2004; Novoa et al., 2001; Novoa et al., 2003). Besides, activated IRE1 can also induce cell apoptosis by associating with tumor necrosis associated factor 2 (TRAF2) and the proapoptotic BCL-2 family members BAX and BAK (Hetz et al., 2006; Ron and Walter, 2007; Urano et al., 2000; Yoneda et al., 2001). Although there are still arguments, it is thought that IRE1 acts to promote cell survival, whereas PERK acts to promote apoptosis (Chang et al., 2018; Lin et al., 2009). Therefore, cell fate is a combined result of responses of the two parallel branches to ER stress.

In addition to intra-branch regulation, previous studies also suggested that there is interbranch regulation between the IRE1 and PERK pathways. XBP1s induces the expression of p58^IPK^ (Kanemoto et al., 2005; Lee et al., 2003), which represses the activation of the PERK branch by inhibiting the kinase activity of PERK (Huber et al., 2013; van Huizen et al., 2003; Yan et al., 2002). Furthermore, a recent study demonstrated that the phosphatase RNA polymerase II-associated protein 2 (RPAP2), which is induced by phosphorylation of eIF2α, dephosphorylates IRE1 (Chang et al., 2018). Therefore, we propose that the IRE1 and PERK branches have mutual inhibition and their interaction regulates cellular response to different ER stress levels.

Here to quantitatively study how they interact in response to different ER stress levels, we constructed a dual-fluorescence reporter to monitor the activity of IRE1 and PERK branch simultaneously in the same cell. Using this system, we quantified the IRE1 and PERK activity over time and over a wide range of ER stresses. We confirmed IRE1 and PERK show mutual inhibition over a wide range of ER stress levels by mediators p58^IPK^ and RPAP2. Furthermore, we found that IRE1 branch responds more sensitively than the PERK branch to low ER stress. This differential sensitivity is dependent on the mutual inhibition by p58^IPK^, rather than the inner-branch inhibition by GADD34. Thus, we proposed that IRE1 and PERK branch coordinate with each other according to the ER stress level via mutual inhibition.

## Results and Discussions

### Construction of the dual-reporter cell system

To simultaneously analyze the dose response of the IRE1 and PERK branches in the same cell, we transformed the HEK 293 cells with both the fluorescence reporter for IRE1 branch activity and fluorescence reporter for PERK branch activity (Walter et al., 2015). The reporter of IRE1 activity is constructed by incorporating a fluorescent protein tagged XBP1 ORF (XBP1-GFP) under the control of an SSFV promoter into the genome of HEK 293 cells. Under ER stress, XBP1-GFP mRNA is spliced by activated IRE1 which has the endonuclease activity. Only this spliced XBP1s-GFP can produce XBP1-GFP fusion protein. The reporter of PERK activity is constructed by incorporating an ATF4 promoter with a CMV promoter and in frame with a fluorescent protein mCherry (Figure 1A). As shown in both fluorescent microcopy and flow cytometry, the IRE1 and PERK branches were clearly activated after the addition of thapsigargin (Tg), an ER stress chemical inducer (Figure 1B-C). The time course of the activation after the treatment of 500 nM Tg showed that the IRE1 branch responded to the stress much faster than the PERK branch. IRE1 activity peaked after 6 hours of Tg treatment, whereas PERK activity increased continuously over time (Figure 1C). Our observation agreed with previous studies, i.e., the IRE1 branch was activated more quickly and attenuated earlier than the PERK branch (Lin et al., 2007; Walter et al., 2015).

**Figure 1.**
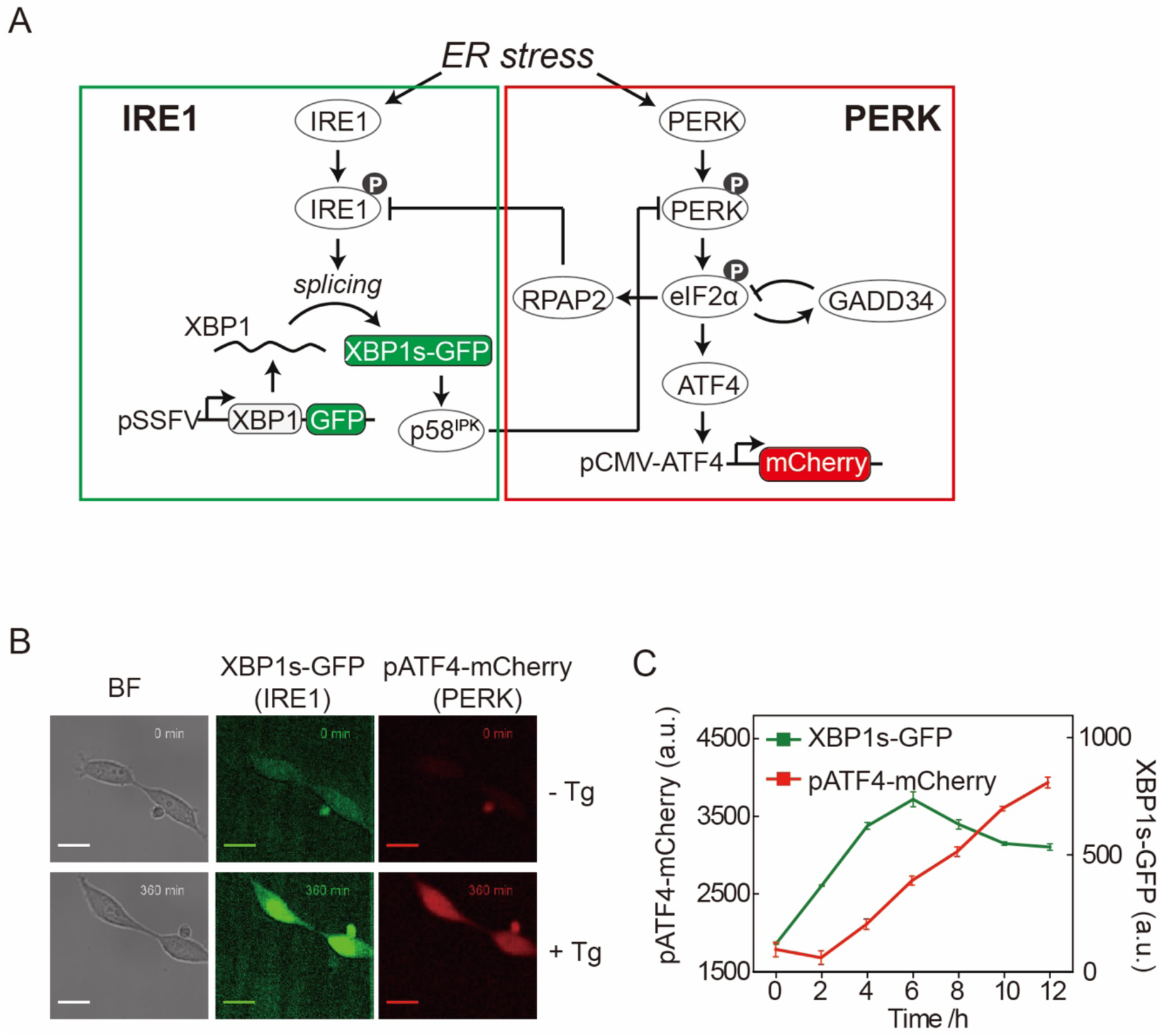
Construction of the dual-reporter system. (A) The design of the dual-reporter cell line that was used to monitor the activity level of the IRE1 and PERK branches by the fluorescence level of GFP and mCherry, respectively. (B) Representative microscopy images of the dual-reporter cell line. T = 0 min (upper panel) and T = 360 min (lower panel) after treatment with 500 nM Tg. Scale bar, 10 μm. (C) The time course of the fluorescence intensities of XBP1s-GFP and pATF4-mCherry after treatment with 500 nM Tg. GFP and mCherry fluorescence were quantified by flow cytometry.

### The IRE1 and PERK branches show mutual inhibition under ER stress

We next investigated how the IRE1 and PERK branches interact with each other upon different ER stress levels by perturbing the potential mediators. We compared the responses of different cell lines at 6 h after exposed to different Tg concentrations. As mentioned above, a key mediator acting downstream of IRE1 branch that represses the PERK activity is p58^IPK^. We knocked out p58^IPK^ in the genome by CRISPR/Cas9 system and constructed a mutant cell line (p58^IPK-/-^) (Figure S1). PERK activity of p58^IPK-/-^ cell line was significantly elevated over the entire dose range (Figure 2A), comparing to the control cell line. Thus, p58^IPK^ showed inhibition on PERK branch activity over a wide range of ER stress levels.

**Figure 2.**
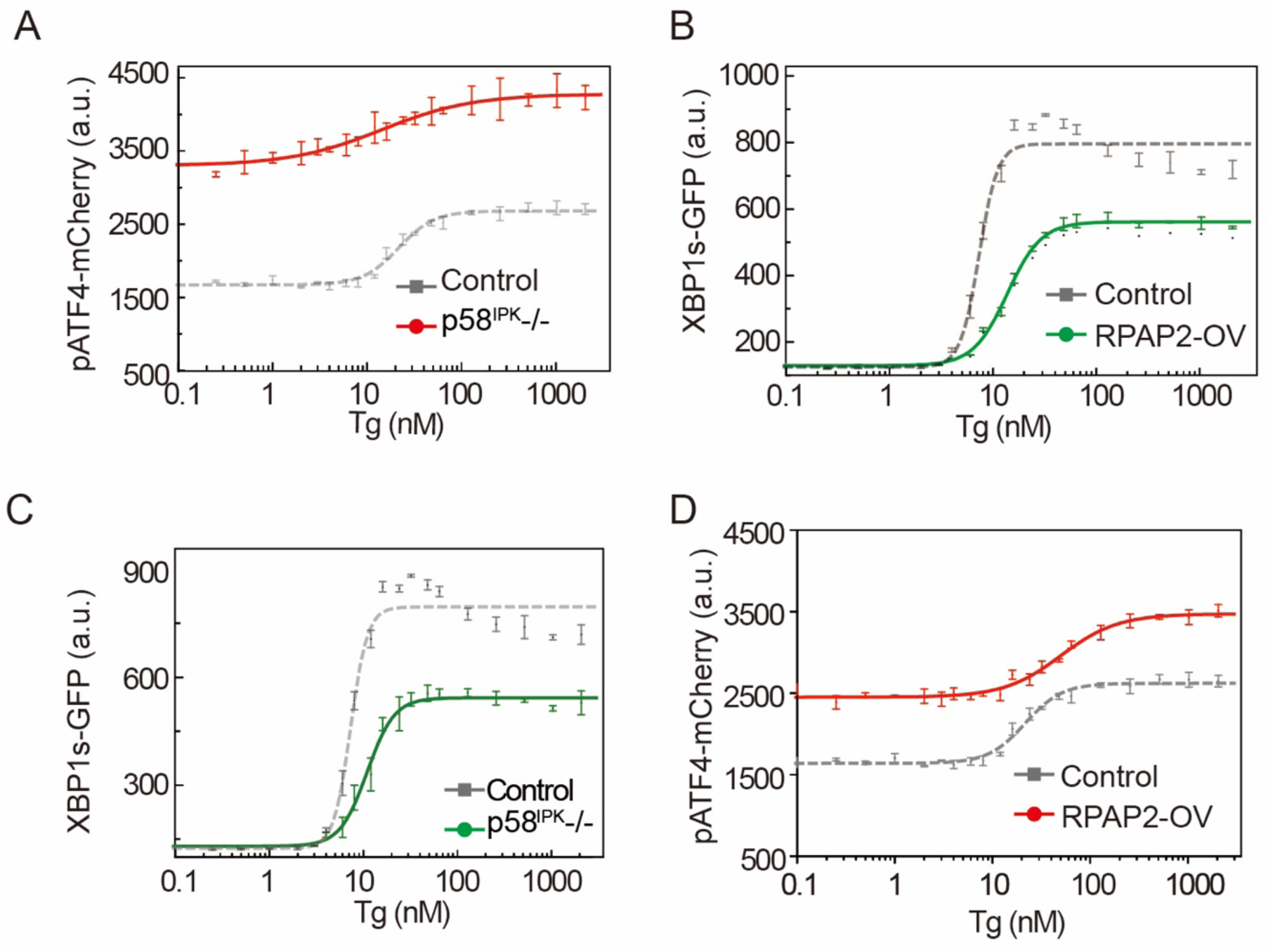
The IRE1 branch and PERK branch have mutual inhibition. (A) pATF4-mCherry dose-response curves of the control (gray) and p58^IPK^-/- (red) cell lines after 6 h of Tg treatment. Knockout of p58^IPK^ downregulated the EC50 of the dose-response curve. Curves, exponential fits. The error bars denote the standard error (n=3). (B) XBP1s-GFP dose-response curves of the control (gray) and RPAP2-OV (green) cell lines after 6 h of Tg treatment. The error bars denote the standard error (n=3). (C) XBP1s-GFP dose-response curves of the control (gray) and p58^IPK^-/- (green) cell lines after 6 h of Tg treatment. The error bars denote the standard error (n=3). (D) pATF4-mCherry dose-response curves of the control (gray) and RPAP2-OV (red) cell lines after 6 h of Tg treatment. The error bars denote the standard error (n=3).

Then we perturbed RPAP2 and tested whether it inhibited IRE1 activity over the entire dose range. As RPAP2 is essential for mammalian cells, we constructed a RPAP2 over-expression cell line (RPAP2-OV) (Figure S2). We found that IRE1 activity in RPAP2-OV cell line was significantly lower than the control cell line when Tg concentration was higher than 6nM. (Figure 2B). RPAP2 overexpression also showed inhibition on IRE1 activity in response to a wide range of ER stress levels, although not under very low ER stress levels.

These results suggested that the inhibition of IRE1 activity by RPAP2 and the inhibition of PERK activity by p58^IPK^ constitutes the mutual inhibition between IRE1 and PERK branch. If so, the perturbation of p58^IPK^ will in turn inhibit IRE1 activity. Indeed, we found that the IRE1 activity in the p58^IPK-/-^ cell line significantly decreased comparing to the control cell line (Figure 2C). Similarly, RPAP2 overexpression also showed increased PERK activity (Figure 2D). Taken together, p58^IPK^ and RPAP2 constitutes the mutual inhibition between IRE1 and PERK branch in a wide range of ER stress levels.

### The interbranch inhibition, rather than the intra-branch inhibition, leads to differential sensitivity of IRE1 and PERK branch

To quantify the response of the signaling pathway to different ER stress levels, we calculated the half maximal effective dose (EC50) for each branch. Notably, the IRE1 activity was slightly attenuated when the Tg reach further higher dosage. This attenuation of IRE1 activity was decreased by perturbation of the regulators of both IRE1 and PERK branch, i.e., p58^IPK^ -/- (Figure 2C), RPAP2-overexpression (Figure 2B) and PERK-/- (Figure S3). We found IRE1 branch was more sensitive than the PERK branch in response to different ER stress levels (Figure 3A). This differential sensitivity was not unique to the time of the Tg treatment. IRE1 branch was sensitive than PERK branch when treated by ER stress for different durations (Figure S4: 4, 8, 10 and 12 h). This differential sensitivity also was not unique to the drugs used to induce ER stress. IRE1 branch was more sensitive than PERK branch in response to other ER-stress-inducing drugs (i.e., tunicamycin (Tm), brefeldin A (BFA)), although the absolute value of EC50_IRE1_ and EC50_PERK_ varied (Figure S5).

**Figure 3.**
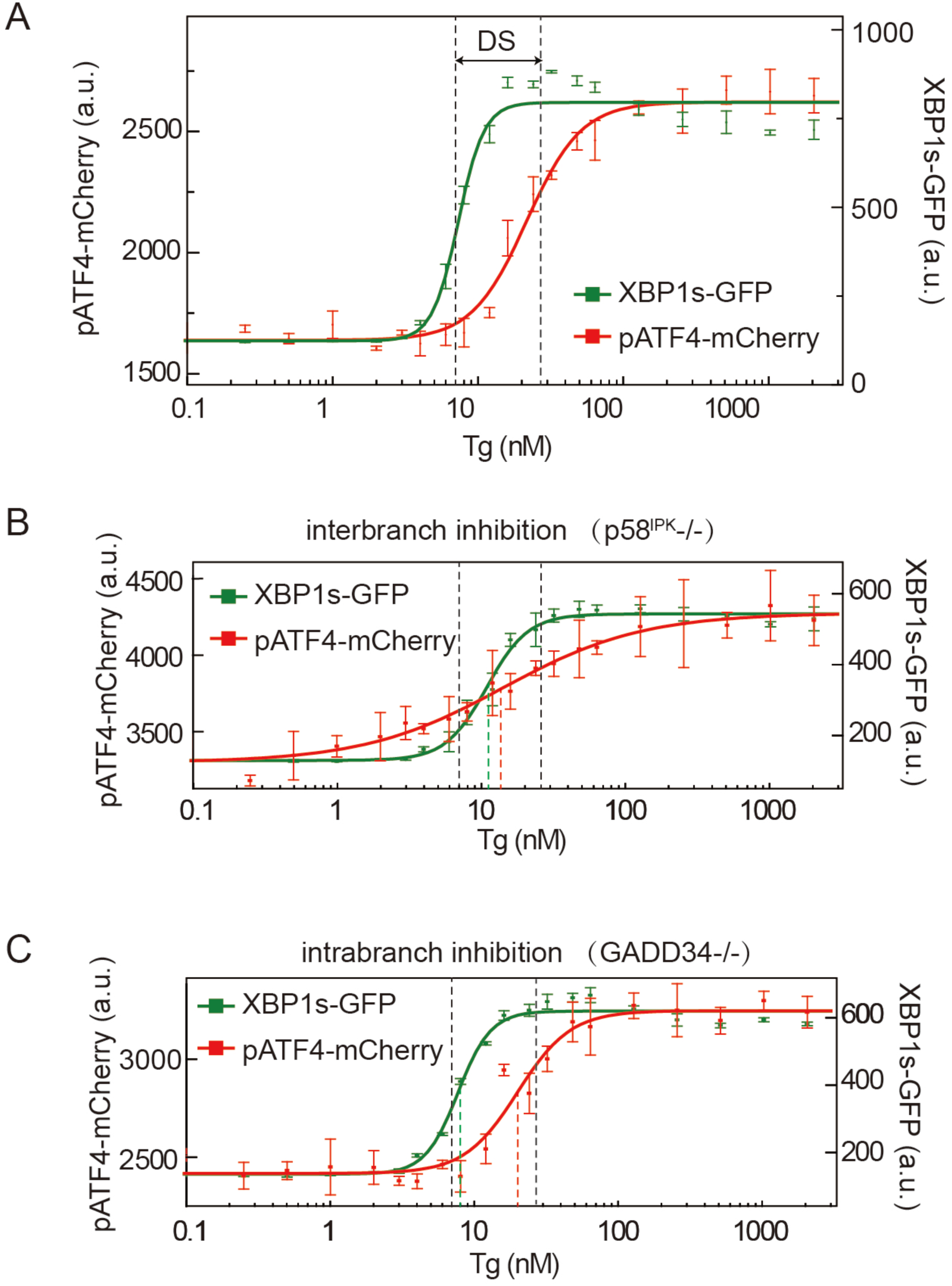
The differential sensitivity is induced by interbranch inhibition, rather than intra-branch inhibition. (A) Dose-response curves for XBP1s-GFP and pATF4-mCherry after 6 h of Tg treatment. GFP and mCherry fluorescence were quantified by flow cytometry. The error bars in the upper panel indicate the standard error of the experimental data (n=3). The black dashed lines show the EC50 for the curves (EC50_IRE1_ = 7 and EC50_PERK_ = 25). The range between the black dashed lines is defined as the DS range. (B) Dose-response curves for XBP1s-GFP and pATF4-mCherry in the p58^IPK^ -/- cell line (EC50_IRE1_ = 11 and EC50_PERK_ = 14). The dashed lines show the EC50 of the indicated curves in the p58^IPK^ -/- cell line. (C) Dose-response curves for XBP1s-GFP and pATF4-mCherry in the GADD34-/- cell line after 6 h of Tg treatment. The dashed lines show the EC50 of the indicated curves in the GADD34-/- cell line (EC50_IRE1_ = 8 and EC50_PERK_ = 20).

To test whether the mutual inhibition affect the differential sensitivity of IRE1 and PERK branch, we quantified the difference of this sensitivity by defining the difference between EC50_IRE1_ and EC50_PERK_ as the differential sensitivity (DS) range (black dashed lines shown in Figure 3A) and compared DS range between different cell lines. In p58^IPK^-/- cell line, DS range was largely decreased (Figure 3B). Next, we investigated whether the intra-branch inhibitor of the PERK branch GADD34 also contributed to inducing the differential sensitivity. We knocked out GADD34 and found that DS range was not obviously altered (Figure 3D and S6). These results suggested that the mutual inhibition has an important role of regulating the activity level of the two branches to different ER stress levels.

Taken together, we proposed a model of how cell integrates parallel signaling pathways to fine-tune its activity level according to the ER stress level. IRE1 and PERK branch mutually inhibit each other’s response to ER stress. By this interbranch inhibition, the IRE1 branch acts more sensitively than PERK branch and the IRE1 branch activation is attenuated at high ER stress. As IRE1 and PERK branch induce different effects on cell fate, this interbranch inhibition may fine-tune the cell fate decision between life and death according to the ER stress level.

## Material and methods

### Cell culture and treatments

The HEK 293 cells were grown in DMEM medium. Drugs used for ER stress treatment were Thapsigargin (T9033, Sigma-Aldrich), tunicamycin (T7765, Sigma-Aldrich) and brefeldin A (B6542, Sigma-Aldrich).

### Cell line construction

A standard protocol of genome editing by Crispr/Cas9 system was used through all the gene knockout process (Ran et al., 2013). For the p58^IPK^ knockout plasmid, 2 guide RNAs (5’-AGGCCGTCGGGGCGGCAGTA-3’ and 5’-TGGCCTCCCAGCGCCGACGG-3’) were used. For the PERK knockout plasmid, 2 guide RNAs (5’-AGGCCGTCGGGGCGGCAGTA-3’ and TGGCCTCCCAGCGCCGACGG) were used. For the GADD34 knockout plasmid, 2 guide RNAs (5’-GGGCGTGGCCGAGATCAGAA-3’ and TGGCATGTATGGTGAGCGAG) were used.

### Fluorescence microscopy

After 2-3 days culture, cells were transferred to a Fibronectin-coated glass-bottom petri dish. After overnight culture, these cells were set for time lapse imaging. Fibronectin (PHE0023, ThermoFisher). All images were captured by a Nikon Observer microscope (Nikon Co., Tokyo, Japan) using a 40 × objective. We used filter sets that were optimized for the detection of GFP and mCherry fluorescent protein and acquired images at 10-minute intervals for 12 h. The exposure time for GFP and mCherry is 50 ms. Cells, objectives and microscope stage were kept at 37 °C and 5% CO_2_ in an environment-control chamber.

### Flow cytometry

Flow cytometry was used for measuring the reporters of IRE1 and PERK branch activity in response to different ER-stress-inducing drugs. Cells in the culture dish were washed twice using 1 × PBS. 4 % paraformaldehyde was used for cell fixation. Green and red lasers were used for measuring the IRE1 branch activity and PERK branch activity, respectively. Cells with no treatment of drugs were measured as experimental control.

## Author contributions

P.W., J.J. and Y.Q. designed the project. Y.Q. and J.J. designed, performed the experiments, and analyzed the data. X.L. wrote computer programs. P.W. and C.T. supervised the whole project. P.W., J.J. and Y.Q. wrote the paper.

---

**Figure S1.**
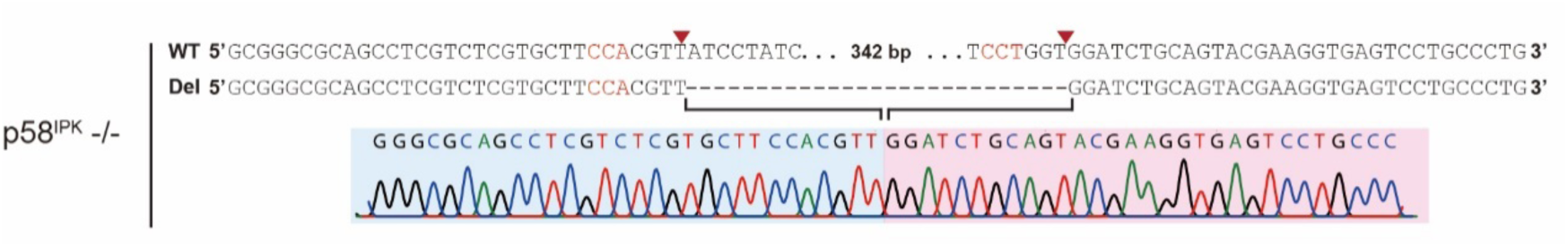
Sanger sequencing traces of the p58^IPK^-/- cell line. Construction of the p58^IPK^-/- cell line using the CRISPR/Cas9 system and a pair of sgRNAs. The sequences of the sgRNAs used to construct the indicated cell lines are shown in Methods. Sanger sequencing traces are shown for the control (upper) and p58^IPK^-/- (lower) cell lines.

**Figure S2.**
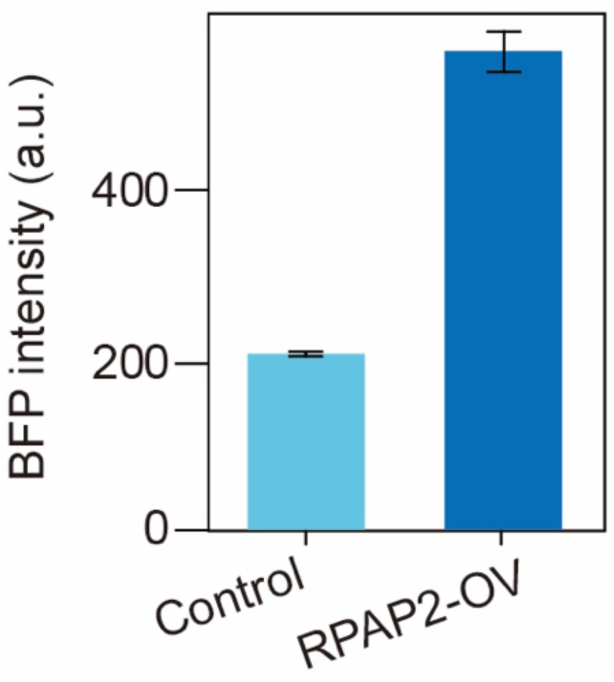
Validation of the RPAP2-OV cell line by FACS. BFP intensity of the control and RPAP2-OV cell lines after treatment with Tg for 6 h. The RPAP2-OV cell line was constructed by transfecting the control cell line with a vector carrying RPAP2-BFP driven by a SFFV promoter. The error bars denote the standard error (n=3).

**Figure S3.**
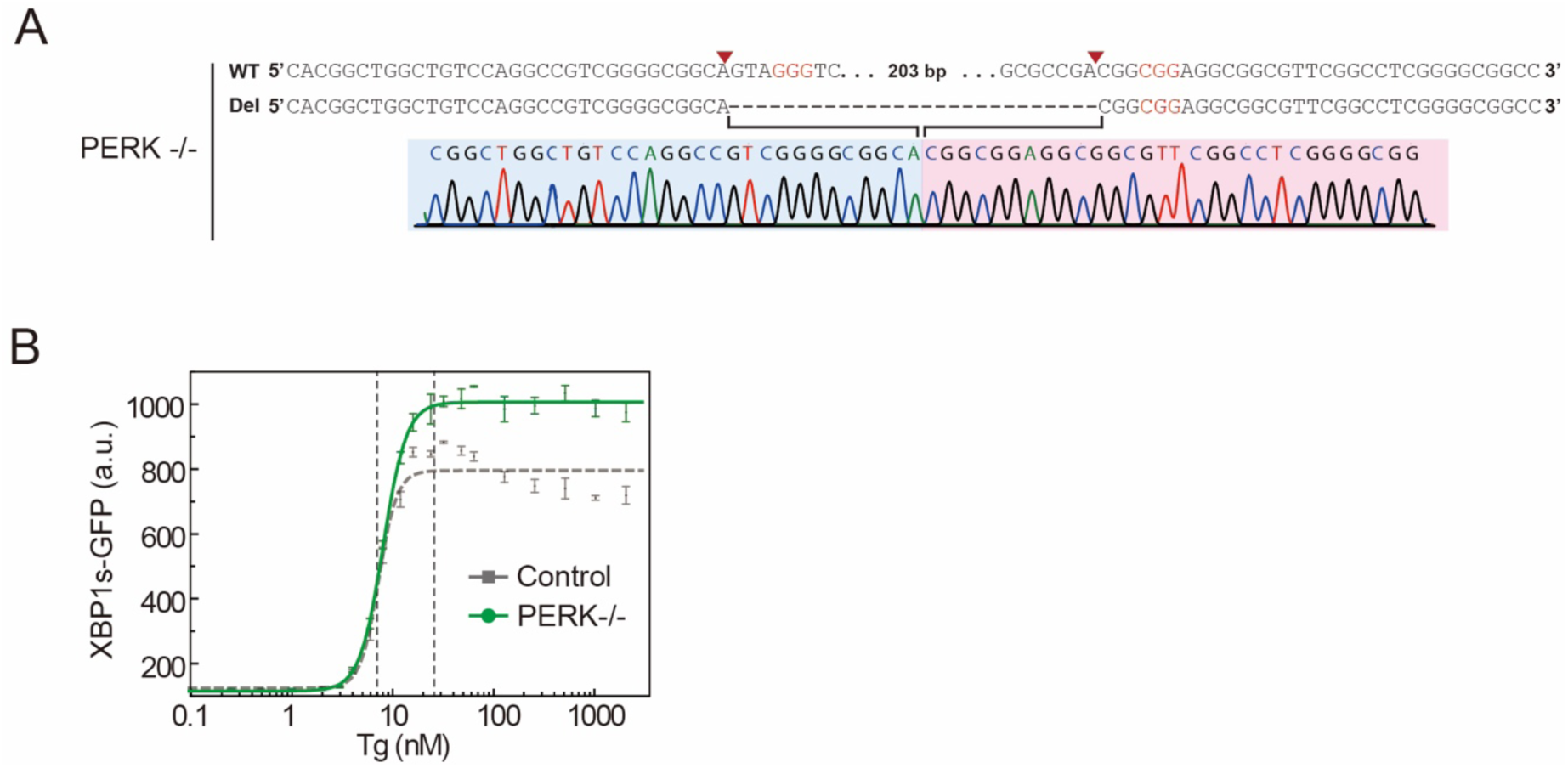
PERK deletion decreased the attenuation of IRE1 activation in response to high Tg concentrations. (A) Construction of the PERK-/- cell line using the CRISPR/Cas9 system and a pair of sgRNAs. Sanger sequencing traces of the control (upper) and PERK-/- (lower) cell lines are shown. (B) XBP1s-GFP intensity in in the control (gray) and PERK-/- (green) cell lines after 6 h of treatment with varying concentrations of Tg.

**Figure S4.**
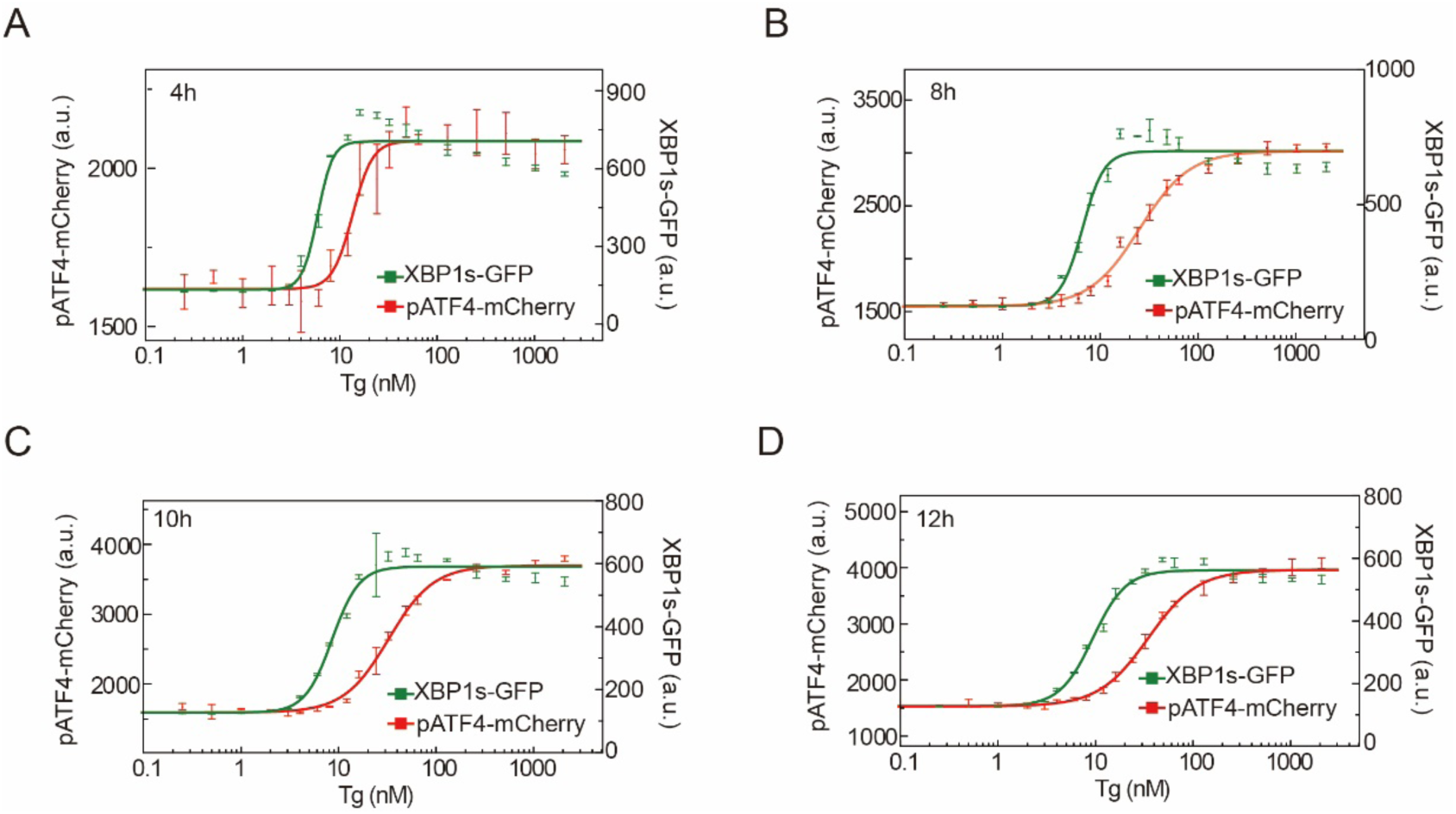
Dose-response curves for XBP1s-GFP and pATF4-mCherry after treatment with Tg for 4, 8, 10 and 12 h. Dose-response curves for XBP1s-GFP and pATF4-mCherry after treatment with Tg for 4 h (A), 8 h (B), 10 h (C) and 12 h (D). The error bars in the upper panel denote the standard error of the experimental data (n=3). (A) EC50_IRE1_ = 6 and EC50_PERK_ = 14; (B) EC50_IRE1_ = 7 and EC50_PERK_ = 25; (C) EC50_IRE1_ = 9 and EC50_PERK_ = 32; and (D) EC50_IRE1_ = 9 and EC50_PERK_ = 35.

**Figure S5.**
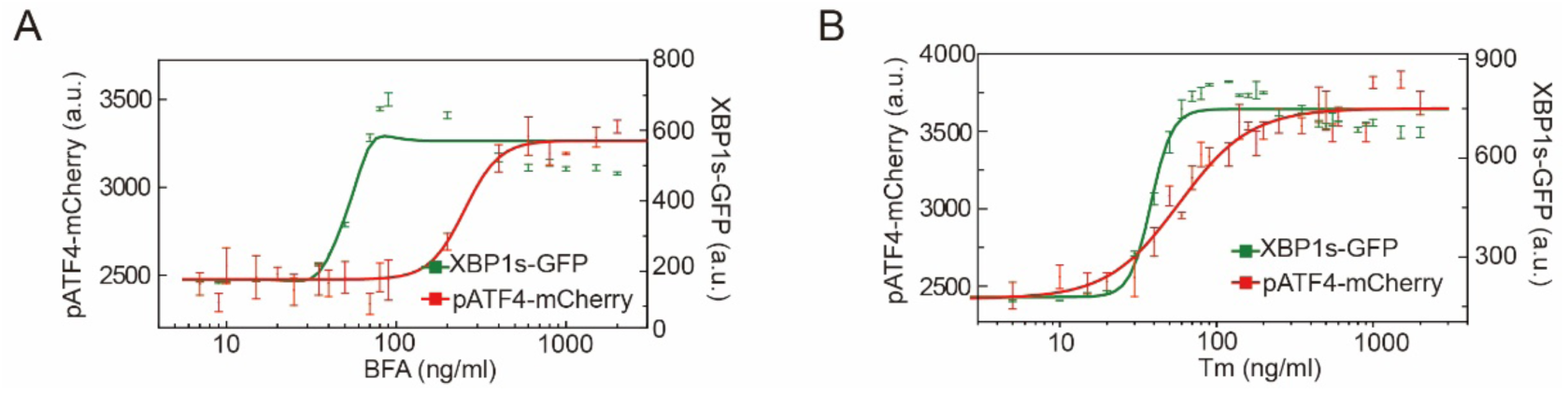
Dose-response curves for XBP1s-GFP and pATF4-mCherry after treatment with BFA and Tm. Dose-response curves for XBP1s-GFP and pATF4-mCherry after 6 h of treatment with BFA (A) and Tm (B). The error bars in the upper panel denote the standard error of the experimental data (n=3). (A) EC50_IRE1_ = 51 and EC50_PERK_ = 252; and (B) EC50_IRE1_ = 38 and EC50_PERK_ = 58.

**Figure S6.**
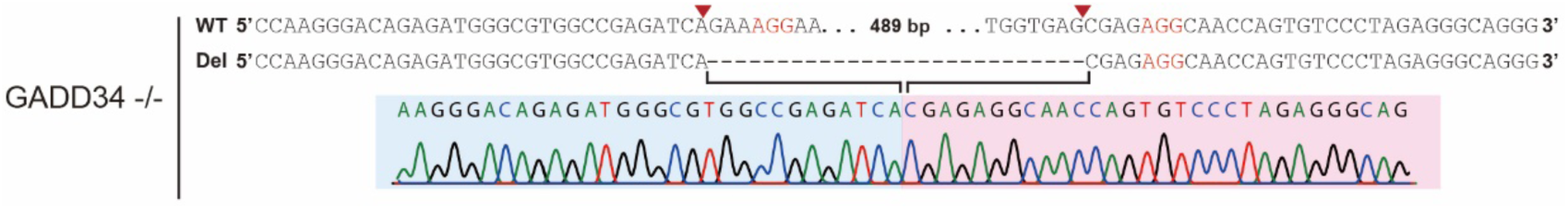
Sanger sequencing traces of the GADD34-/- cell line. Construction of the GADD34-/- cell line using the CRISPR/Cas9 system and a pair of sgRNAs. Sanger sequencing traces of the control (upper) and GADD34-/- (lower) cell lines are shown.

## References

Calfon, M., Zeng, H., Urano, F., Till, J.H., Hubbard, S.R., Harding, H.P., Clark, S.G., and Ron, D. (2002). IRE1 couples endoplasmic reticulum load to secretory capacity by processing the XBP-1 mRNA. Nature 415, 92–96.

Chang, T.K., Lawrence, D.A., Lu, M., Tan, J., Harnoss, J.M., Marsters, S.A., Liu, P., Sandoval, W., Martin, S.E., and Ashkenazi, A. (2018). Coordination between Two Branches of the Unfolded Protein Response Determines Apoptotic Cell Fate. Mol Cell 71, 629–636 e625.

Harding, H.P., Novoa, I., Zhang, Y., Zeng, H., Wek, R., Schapira, M., and Ron, D. (2000). Regulated translation initiation controls stress-induced gene expression in mammalian cells. Mol Cell 6, 1099–1108.

Harding, H.P., Zhang, Y., and Ron, D. (1999). Protein translation and folding are coupled by an endoplasmic-reticulum-resident kinase. Nature 397, 271–274.

Hetz, C., Bernasconi, P., Fisher, J., Lee, A.H., Bassik, M.C., Antonsson, B., Brandt, G.S., Iwakoshi, N.N., Schinzel, A., Glimcher, L.H., et al. (2006). Proapoptotic BAX and BAK modulate the unfolded protein response by a direct interaction with IRE1alpha. Science 312, 572–576.

Hollien, J., Lin, J.H., Li, H., Stevens, N., Walter, P., and Weissman, J.S. (2009). Regulated Ire1-dependent decay of messenger RNAs in mammalian cells. J Cell Biol 186, 323–331.

Hollien, J., and Weissman, J.S. (2006). Decay of endoplasmic reticulum-localized mRNAs during the unfolded protein response. Science 313, 104–107.

Huber, A.L., Lebeau, J., Guillaumot, P., Petrilli, V., Malek, M., Chilloux, J., Fauvet, F., Payen, L., Kfoury, A., Renno, T., et al. (2013). p58(IPK)-mediated attenuation of the proapoptotic PERK-CHOP pathway allows malignant progression upon low glucose. Mol Cell 49, 1049–1059.

Jonikas, M.C., Collins, S.R., Denic, V., Oh, E., Quan, E.M., Schmid, V., Weibezahn, J., Schwappach, B., Walter, P., Weissman, J.S., et al. (2009). Comprehensive characterization of genes required for protein folding in the endoplasmic reticulum. Science 323, 1693–1697.

Kanemoto, S., Kondo, S., Ogata, M., Murakami, T., Urano, F., and Imaizumi, K. (2005). XBP1 activates the transcription of its target genes via an ACGT core sequence under ER stress. Biochem Biophys Res Commun 331, 1146–1153.

Lee, A.H., Iwakoshi, N.N., and Glimcher, L.H. (2003). XBP-1 regulates a subset of endoplasmic reticulum resident chaperone genes in the unfolded protein response. Mol Cell Biol 23, 7448–7459.

Lin, J.H., Li, H., Yasumura, D., Cohen, H.R., Zhang, C., Panning, B., Shokat, K.M., Lavail, M.M., and Walter, P. (2007). IRE1 signaling affects cell fate during the unfolded protein response. Science 318, 944–949.

Lin, J.H., Li, H., Zhang, Y., Ron, D., and Walter, P. (2009). Divergent effects of PERK and IRE1 signaling on cell viability. PLoS One 4, e4170.

Lu, P.D., Harding, H.P., and Ron, D. (2004). Translation reinitiation at alternative open reading frames regulates gene expression in an integrated stress response. Journal of Cell Biology 167, 27–33.

Ma, Y.J., and Hendershot, L.M. (2003). Delineation of a negative feedback regulatory loop that controls protein translation during endoplasmic reticulum stress. Journal of Biological Chemistry 278, 34864–34873.

Marciniak, S.J., Yun, C.Y., Oyadomari, S., Novoa, I., Zhang, Y.H., Jungreis, R., Nagata, K., Harding, H.P., and Ron, D. (2004). CHOP induces death by promoting protein synthesis and oxidation in the stressed endoplasmic reticulum. Gene Dev 18, 3066–3077.

Novoa, I., Zeng, H.Q., Harding, H.P., and Ron, D. (2001). Feedback inhibition of the unfolded protein response by GADD34-mediated dephosphorylation of eIF2 alpha. Journal of Cell Biology 153, 1011–1021.

Novoa, I., Zhang, Y.H., Zeng, H.Q., Jungreis, R., Harding, H.P., and Ron, D. (2003). Stress-induced gene expression requires programmed recovery from translational repression. Embo J 22, 1180–1187.

Puthalakath, H., O’Reilly, L.A., Gunn, P., Lee, L., Kelly, P.N., Huntington, N.D., Hughes, P.D., Michalak, E.M., McKimm-Breschkin, J., Motoyama, N., et al. (2007). ER stress triggers apoptosis by activating BH3-only protein Bim. Cell 129, 1337–1349.

Ran, F.A., Hsu, P.D., Wright, J., Agarwala, V., Scott, D.A., and Zhang, F. (2013). Genome engineering using the CRISPR-Cas9 system. Nat Protoc 8, 2281–2308.

Ron, D., and Walter, P. (2007). Signal integration in the endoplasmic reticulum unfolded protein response. Nat Rev Mol Cell Biol 8, 519–529.

Urano, F., Wang, X., Bertolotti, A., Zhang, Y., Chung, P., Harding, H.P., and Ron, D. (2000). Coupling of stress in the ER to activation of JNK protein kinases by transmembrane protein kinase IRE1. Science 287, 664–666.

van Huizen, R., Martindale, J.L., Gorospe, M., and Holbrook, N.J. (2003). P58IPK, a novel endoplasmic reticulum stress-inducible protein and potential negative regulator of eIF2alpha signaling. J Biol Chem 278, 15558–15564.

Walter, F., Schmid, J., Dussmann, H., Concannon, C.G., and Prehn, J.H. (2015). Imaging of single cell responses to ER stress indicates that the relative dynamics of IRE1/XBP1 and PERK/ATF4 signalling rather than a switch between signalling branches determine cell survival. Cell Death Differ 22, 1502–1516.

Walter, P., and Ron, D. (2011). The unfolded protein response: from stress pathway to homeostatic regulation. Science 334, 1081–1086.

Yan, W., Frank, C.L., Korth, M.J., Sopher, B.L., Novoa, I., Ron, D., and Katze, M.G. (2002). Control of PERK eIF2alpha kinase activity by the endoplasmic reticulum stress-induced molecular chaperone P58IPK. Proc Natl Acad Sci U S A 99, 15920–15925.

Yoneda, T., Imaizumi, K., Oono, K., Yui, D., Gomi, F., Katayama, T., and Tohyama, M. (2001). Activation of caspase-12, an endoplastic reticulum (ER) resident caspase, through tumor necrosis factor receptor-associated factor 2-dependent mechanism in response to the ER stress. J Biol Chem 276, 13935–13940.

